# 16GT: a fast and sensitive variant caller using a 16-genotype probabilistic model

**DOI:** 10.1101/111393

**Authors:** Ruibang Luo, Michael C. Schatz, Steven L. Salzberg

## Abstract

**Summary:** 16GT is a variant caller for Illumina WGS and WES germline data. It uses a new 16-genotype probabilistic model to unify SNP and indel calling in a single variant calling algorithm. In benchmark comparisons with five other widely used variant callers on a modern 36-core server, 16GT ran faster and demonstrated improved sensitivity in calling SNPs, and it provided comparable sensitivity and accuracy in calling indels as compared to the GATK HaplotypeCaller.

**Availability and implementation:** https://github.com/aquaskyline/16GT

**Supplementary information:** Supplementary tables and notes are available at *Bioinformatics* online.

## 1 Introduction

Single nucleotide polymorphisms (SNPs) and insertions and deletions (indels) that occur at a specific genome position are interdependent; i.e., evidence that elevates the probability of one variant type should decrease the probability of other possible vaemailriant types, and the probability of all possible alleles should sum up to 1. However, widely-used tools such as GATK’s UnifiedGenotyper (McKenna, et al., 2010) and SAMtools (Li, et al., 2009) use separate models for SNP and indel detection. The model for SNP calling in these two tools is nearly identical: both assume all variants are biallelic and use a probabilistic model allowing for 10 genotypes (AA, AC, AG, AT, CC, CG, CT, GG, GT, TT). For indel calling, the GATK UnifiedGenotyper uses a model from Dindel’s model (Albers, et al., 2011), while SAMtools’ model is from BAQ (Li, 2011).

## 2 Methods

In order to detect SNPs and indels with a unified approach, we developed a new 16-genotype probabilistic model and its implementation named 16GT. The idea was firstly introduced by (Luo, et al., 2014), 16GT implements the idea using an improved model. Using X and Y to denote the indels with the highest (X) and second highest (Y) support, we add 6 new genotypes (AX, CX, GX, TX, XX and XY) to the traditional 10-genotype probabilistic model. The six new genotypes include: 1) one homozygous indel (XX); 2) one reference allele plus one heterozygous indel (AX, CX, GX, TX); 3) one heterozygous SNP plus one heterozygous indel (AX, CX, GX, TX, reusing the genotypes in 2); and 4) two heterozygous indels (XY). We exclude the 5 possible combinations AY, CY, GY, TY, YY because X has higher support than Y. By unifying SNP and indel calling in a single variant calling algorithm, 16GT not only runs 4 times faster, but also demonstrates improved sensitivity in calling SNPs and comparable sensitivity in calling indels to the GATK HaplotypeCaller.

Posterior probabilities of these 16 genotypes are calculated using a Bayesian model *P*(*L\F*)∝*P*(*F\L*)*P*(*L*), where *L* is an assumed genotype. *F* refers to the observation of the 6 alleles (A, C, G, T, X, Y) at a given genome position. *P*(*L*) is the prior probability of the genotype, *P*(*F*\*L*) is the likelihood of the observed genotype. and *P*(*L*\*F*) is the posterior probability of the genotype. The resulting genotype *L_max_* is assigned to the genotype with the highest posterior probability.

### 2.1 Calculate *P*(*F|L*)

To test how well an observation fits the expectation of different genotypes, we use a two-tailed Fisher’s Exact Test *P* and use the resulting *p*-value as the goodness of fit. When calculating the likelihood of a homozygous genotype, ideally we expect 100% single allele support from the observation. For example, consider genotype ‘AA’:

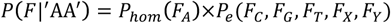

For a heterozygous genotype, 50% support is expected for each allele in the genotype, for example consider ‘CG’:

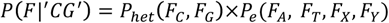

where

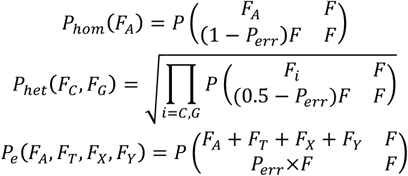

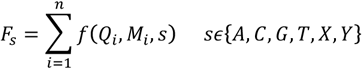

where *s* is the allele type, *n* is the number of bases supporting allele *s*, *Q_i_* is the base quality, and *M_i_* is the mapping quality. *f* is a function describing how *s*, *Q_i_* and *M_i_* change the observation:

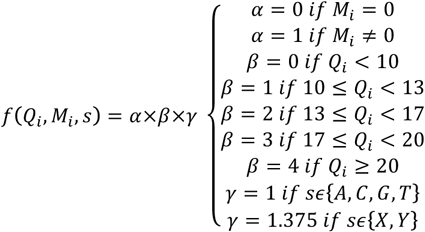

The possible reasons for an observation that does not match the reference genome are: 1) a true variant; 2) error generated in library construction; 3) base calling error; 4) mapping error; and 5) error in the reference genome. Reasons 3 and 4 are explicitly captured in our model. For reasons 2 and 5, we include two error probabilities, *P_s_* for SNP error and *P_d_* for indel error. We define *P_err_*=*P_s_*+*P_d_*, where *P_s_* and *P_d_* are set to 0.01 and 0.005, respectively. These two values were set empirically based on the observation that SNP errors are more common than indel errors in library construction and in the reference genome.

In addition, most short read aligners use a dynamic programming algorithm to enable gapped alignment, using a scoring scheme that usually penalizes gap opening and extension more than mismatch. Consequently, authentic gaps that occur at an end of a read are more likely to be substituted by a set of false SNPs or alternatively to get trimmed or clipped. Thus, we applied a coefficient γ to weight indel observation more than SNP to increase the sensitivity on indels.

### 2.2 Calculate *P*(*L*)

Given 1) a known rate of single nucleotide differences between two unrelated haplotypes; 2) a known rate of single indel differences between two unrelated haplotypes; and 3) a known T?ransi?tions to T?ransv?ersions rtio (Ti/Tv), the 16GT model’s prior probabilities are as calculated as shown in **Supplementary Table 1.**

## 3 Results

We benchmarked 16GT with GATK UnifiedGenotyper, GATK HaplotypeCaller (McKenna, et al., 2010), Freebayes (Garrison and Marth, 2012), Fermikit (Li, 2015) and ISAAC (Raczy, et al., 2013) using a set of very high-confidence variants developed by the Ge-nome-in-a-bottle (GIAB) project for genome NA12878 (Zook, et al., 2014) **(Supp. Note)**. The results are shown in Table 1. For SNPs, 16GT produced the fewest false negative calls; i.e., it missed the smallest number of true SNPs, and 79% of 16GT’s false positive calls were also reported by dbSNP version 138, which is highest among other callers. However, we should point out that the GIAB variant set is biased towards GATK because it was primarily derived from GATK-based analyses (Chiang, et al., 2015). As a less-biased test, we therefore assessed the false positive calls against a set of unbiased calls made by the Illumina Omni 2.5 SNP array **(Supp. Note)**. Among the 5,346 false positive calls for 16GT, 20 were covered by Omni 2.5 and all 20 had the correct genotype, which suggests a considerable number of the "false positive calls” are actually correct. For indels, 16GT produced more false negative calls than HaplotypeCaller, but less than half as many false negative calls as UnifiedGenotyper. 65% of 16GT’s false positive indels were covered by dbSNP. Among the 1,462 false positive indels, 981 (67%) of them meet all three of the following criteria: 1) at least three reads supporting the variant; 2) at least one read supporting both the positive and negative strands, and; 3) in over 80% of the reads that support the variant, there exists no other variant in its flanking 10bp. This suggests that some of these “false positive calls” might be correct and require experimental validation to confirm. Supplementary Figure 1 shows three examples where the putative false positive from 16GT is likely to be correct.

**Table 1:** Benchmark between 16GT and five other variant callers on a dataset from the Genome in a Bottle project consisting of 787M read pairs (53-fold) from genome NA12878. UG: GATK UnifiedGenotyper; HC: GATK HaplotypeCaller. FP: false positive, FN: false negative.

| Caller | Time<br>(minutes w/<br>36 cores) | SNP |  |  |  |  | Indel |  |  |  |
| --- | --- | --- | --- | --- | --- | --- | --- | --- | --- | --- |
|  |  | FP |  |  |  | FN | FP |  |  | FN |
|  |  | Total | dbSNP138 | dbSNP138<br>% | TP in<br>Omni 2.5 |  | Total | dbSNP138 | dbSNP138<br>% |  |
| 16GT | 121 | 5,346 | 4,220 | 79% | 20/20 | 918 | 1,462 | 944 | 65% | 3,180 |
| UG | 29 | 1,639 | 563 | 34% | 15/15 | 8,489 | 624 | 546 | 88% | 6,890 |
| HC | 539 | 419 | 143 | 34% | 4/4 | 10,413 | 1,232 | 726 | 59% | 2,285 |
| Freebayes | 52 | 724 | 353 | 49% | 11/14 | 8,584 | 559 | 0 | 0% | 8,224 |
| Fermikit | 45 | 2,036 | 509 | 25% | 9/9 | 96,425 | 1,996 | 1,076 | 54% | 8,813 |
| ISAAC | 63 | 1,115 | 586 | 53% | 15/15 | 4,659 | 1,239 | 710 | 57% | 12,087 |

## 4 Conclusion

Compared with local assembly based variant callers, 16GT provides better sensitivity in SNP calling and comparable sensitivity in indel calling. In the future, we will improve 16GT to support somatic variant detection and species with more than two haplotypes.

## Funding and Acknowledgement

This work has been supported by the U.S. National Institutes of Health under grants R01-HL129239 and R01-HG006677. We thank UEC for providing the bam2snapshot function.

